# A Moment-Based Maximum Entropy Model for Fitting Higher-Order Interactions in Neural Data

**DOI:** 10.1101/353656

**Authors:** N. Alex Cayco-Gajic, Joel Zylberberg, Eric Shea-Brown

## Abstract

Correlations in neural activity have been demonstrated to have profound consequences for sensory encoding. To understand how neural populations represent stimulus information, it is therefore necessary to model how pairwise and higher-order spiking correlations between neurons contribute to the collective structure of population-wide spiking patterns. Maximum entropy models are an increasingly popular method for capturing collective neural activity by including successively higher-order interaction terms. However, incorporating higher-order interactions in these models is difficult in practice due to two factors. First, the number of parameters exponentially increases as higher orders are added. Second, because triplet (and higher) spiking events occur infrequently, estimates of higher-order statistics may be contaminated by sampling noise. To address this, we extend previous work on the Reliable Interaction class of models [1] to develop a normalized variant that adaptively identifies the specific pairwise and higher-order moments that can be estimated from a given dataset for a specified confidence level. The resulting “Reliable Moment” model is able to capture cortical-like distributions of population spiking patterns. Finally, we show that, compared with the Reliable Interaction model, the Reliable Moment model infers fewer strong spurious higher-order interactions and is better able to predict the frequencies of previously unobserved spiking patterns.

## 1. Introduction

An essential step in understanding neural coding is the characterization of the correlated structure of neural activity. Over the past two decades, much theoretical work has clarified the strong impact that correlated variability between pairs of neurons can have on the amount of information that can be encoded in neural circuits [2–7]. Beyond pairs, recent experimental studies have shown evidence of *higher-order* correlations in cortical [8–12] and retinal [1,13] population activity. Depending on their stimulus-dependent structure, these higher-order correlations could also have a strong impact on population coding [14,15]. Moreover, capturing higher-order correlations in neural spiking may be important for identifying functional networks in neural circuits [16], or for characterizing their collective statistical activity [17]. Therefore, to incorporate higher-order spiking statistics into an information theoretic framework, we require flexible modeling tools that can capture the coordinated spiking of arbitrary orders within neural populations.

Maximum entropy models are an increasingly common tool for fitting and analyzing neural population spiking patterns. Intuitively, maximum entropy models fit certain specified features (e.g., firing rates, correlations between cells) while making minimal additional assumptions about the population structure [18]. Several variants of the maximum entropy model have been used to fit the collective activity of spiking patterns in neural data [5,12,13,16,19,20]. However, it is still unclear how to efficiently incorporate higher-order features into maximum entropy models for two reasons. First, the number of parameters (and hence the computational expense of model fitting) increases exponentially as higher-order features are incorporated. Second, because higher-order synchronous spiking occurs infrequently, empirical estimates tend to be noisy; therefore, massive amounts of data may be necessary to create a model with higher-order interactions that can generalize to held-out data. These issues have been addressed by the Reliable Interaction model [1], which uses a maximum entropy inspired model to fit a sparse network of features based on the most “reliable” (i.e., high-frequency) spiking patterns within their data. This approach is extremely efficient numerically and reproduces the frequencies of the most commonly occurring patterns with high accuracy. However, because the model is not a normalized probability distribution, it cannot be used to calculate information theoretic quantities such as the Kullback−Leibler divergence or mutual information.

To address these challenges, we introduce an adaptive maximum entropy model that identifies and fits spiking interactions of all orders, based on the criterion that they can be accurately estimated from the data for a specified confidence level. Towards this end, we adapt the Reliable Interaction model by making two small but critical modifications in the fitting procedure and fitting criterion; these modifications normalize the model, allowing information theoretic quantities to be calculated. The resulting model is able to fit cortical-like distributions of spiking patterns with dense higher-order statistics. Finally, we show that these modifications have two further important consequences: they reduce spurious higher-order interactions, and improve the model’s ability to predict the frequencies of previously unseen spiking patterns.

## 2. Results

### 2.1. The Reliable Moment Model

To analyze population-level activity in neural recordings, it is often necessary to first model the distribution of spiking patterns. Certain spiking features of neural population activity are likely to be more relevant for modeling than others: for example, each neuron’s firing rate and the correlations between pairs of neurons. In general, there may be an infinite family of models that fit these key features in the data, making any particular choice seem potentially arbitrary. One approach is to take the distribution that captures the identified statistical features while making the fewest additional assumptions on the structure of the data. Mathematically, this is equivalent to matching the average values of the features observed in the data while maximizing the statistical entropy [21]. The resulting distribution is called the maximum entropy model and can be derived analytically via Lagrange multipliers [18], resulting in the following probability:

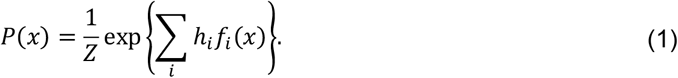

Here, *x* represents a binary spiking pattern across the population in a small time bin (i.e., *x*_*i*_ = 1 if neuron *i* spiked in that time bin, otherwise *x*_*i*_ = 0), *f*_*i*_(*x*) are the chosen spiking features, and h_i_ are interaction parameters that are fitted to match the average 〈*f*_*i*_(*x*)〉 to the values observed in the data. *Z* is a normalizing factor, also called the partition function.

The quality of fit of a maximum entropy model relies critically on which features are included. Traditionally, first-order (i.e., firing rate) and second-order features (correlations) are chosen [5] to isolate the effect of pairwise correlations on population activity patterns. However, this may miss important information about higher-order dependencies within the data. In principle, the pairwise maximum entropy model can be generalized by fitting features of up to kth order; but this becomes computationally expensive for large datasets as the number of parameters grows as O(N^k^). Moreover, higher-order features are more susceptible to overfitting, because they represent spiking features that occur less frequently in the data (and consequently have noisy empirical estimates). An alternative is to incorporate a limited subset of predetermined phenomenological features that increase the predictive power of the model, such as the spike count distribution [13] or frequency of the quiescent state [12]. While these models have been able to capture the collective activity of populations of neurons (e.g., to determine whether neural activity operates at a critical point [17]), they are not able to dissect how the functional connectivity between specific subgroups of neurons contributes to the population level activity.

To address these challenges, a method is needed for data-driven adaptive identification of relevant spiking features of all orders. The Reliable Interaction (RI) model [1] has previously been used to fit sparse networks of pairwise and higher-order interactions to retinal populations. The RI model fits only the features corresponding to spiking patterns whose observed frequencies are larger than an arbitrary threshold. For example, in a 10-cell population, the fourth-order feature *f*_*i*_(*x*) = *x*_1_*x*_3_*x*_5_*x*_9_ would be fitted only if the spiking pattern *x* = 1010100010 occurs with frequency above this threshold. Once these features have been identified, the RI model uses an algebraic approximation for rapid parameter fitting by first calculating the partition function Z as the inverse of the frequency of the silent state: *Z* = *P*(00 …0)^−1^. Subsequently, the interaction parameters can be estimated recursively from the observed frequencies and *Z*. However, while the RI model has been shown to be able to accurately fit the frequencies of spiking patterns, its fitting procedure does not generate a normalized probability distribution (as originally discussed in [1]; see Appendix A for an intuitive example). This limits certain applications of the model: for example, information theoretic measures such as the Kullback−Leibler divergence and mutual information cannot be calculated. Another limitation (demonstrated below and in Appendix A) is that the RI model often cannot predict the frequencies of rarely occurring spiking patterns.

We propose the Reliable Moment (RM) model, an adaptation of the RI model that makes two key modifications in the fitting procedure and fitting criterion. First, we take advantage of a recently developed method for rapid parameter estimation: Minimum Probability Flow (MPF) learning [22]. While still substantially slower than the algebraic method employed in [1] (which is essentially instantaneous), using a parameter estimation method such as MPF guarantees a probability distribution that, in theory, can be readily normalized. In practice, calculating the partition function (*Z* in Equation (1)) may be computationally expensive, as it requires summing 2^N^ probabilities. In this case, the partition function can be quickly estimated using other techniques, such as the Good−Turing estimate [23] (see Methods). As we shall see below, attempting to apply these approaches to the RI model strongly disrupts its predictions.

Second, instead of fitting the features corresponding to the most commonly occurring spiking patterns, we fit the features corresponding to the largest moments. Taking the previous example, feature *f*_*i*_(*x*) = *x*_1_*x*_3_*x*_5_*x*_9_ would be fitted only if the moment 〈*x*_1_*x*_3_*x*_5_*x*_9_〉 is greater than some threshold. As in the RI model, the threshold parameter *p*_*min*_ implicitly determines the number of fitted features. For binary systems, the uncentered moment of a subset of neurons is equal to the marginal probability of those neurons spiking, so that the previous condition is equivalent to:

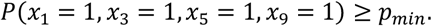

The choice of *p*_*min*_ can be made less arbitrary by choosing its value to bound the 95% confidence interval of the relative error in the sample moments (with some minimal assumptions; [14]):

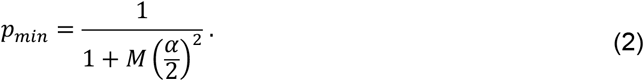

*M* is the number of samples and *α* is the maximum desired relative error. In this way, the RM model can adaptively identify which moments within a specific dataset are large enough to be accurately estimated by the sample frequency.

Unlike the spiking pattern frequencies used in the RI model, these marginal probabilities satisfy an important hierarchy: the moment of any set of neurons is necessarily bounded by the moment of any subset of those neurons; e.g.,:

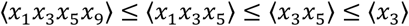

This means that for every higher-order interaction fitted by the RM model, all of its corresponding lower-order interactions are automatically fitted as well. Although this may seem to be a minor change from the RI model, we will demonstrate the significance of this change with the following toy model (we later consider larger and more realistic models, i.e., Figs. 2-5).

### 2.2. Illustration with a Toy Example

Consider *N* = 3 homogeneous neurons with only first and second-order interactions:

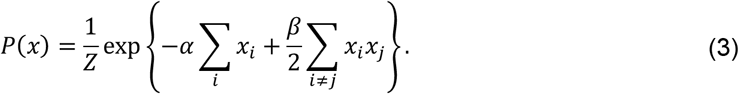

The probability of each pattern can be found analytically:

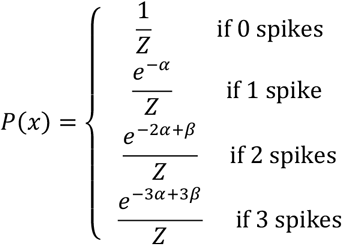

where *Z* = 1 + 3*e*^−*α*^ + 3*e*^−2*α*+*β*^ + *e*^−3*α*+3*β*^. In particular, for *α* = 1, *β* = 1.2:

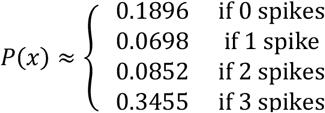

To gain intuition on the fundamental differences between the RM and RI models, we will take the “best-case” scenario for the model fits; i.e., assuming infinite data and infinite fitting time. This eliminates any error due to statistical sampling or parameter fitting for this toy example. We will first see that the difference in fitting criterion can lead the RI model to identify spurious higher-order interactions. This can be seen by setting the threshold at *p*_*min*_ = 0.1. Then, the RI model will only identify the spiking patterns *x* = 000 and 111 as reliable, resulting in the following:

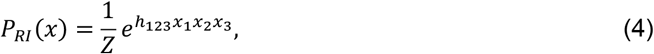

where *h*_123_ = *log*(*Z* * *P*(111)) = 0.6. While the ground truth distribution only contains first- and second-order interactions, the RI fitting procedure mistakenly infers a pure triplet model. This happens because the RI model criterion for selection is based on the frequencies of spiking patterns, which (unlike the moments) do not necessarily follow a natural hierarchy. In contrast, because it relies on the frequency of the marginal probabilities, the RM model identifies all first, second, and third order interaction parameters:

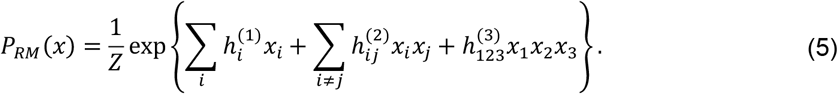

This demonstrates that the RM model cannot infer higher-order interactions without also fitting the corresponding lower-order interactions.

Second, the RI model can fail to predict the frequencies of rare spiking patterns; i.e., those that were not selected as reliable by the model. To see this, consider that the RI model estimates the partition function as *Z* = *P*(000)^−1^. While this gives an accurate estimate of the partition function of the true underlying distribution (in this example, the pairwise model; Equation (3)), it may be a poor estimate of the partition function for the model with interactions inferred by the RI fitting criterion (i.e., the pure triplet model). This mismatch between model form and the estimated partition function is the reason the model cannot be normalized. Because the estimated *Z* is also used to determine the interaction parameters, the RI model frequencies match the true probabilities of the spiking patterns that are used for fitting (i.e., the most common or reliable patterns), but is inaccurate for patterns that are below the threshold frequency (Figure 1). However, naïve renormalization of the model would make all of the probabilities inaccurate.

**Figure 1.**
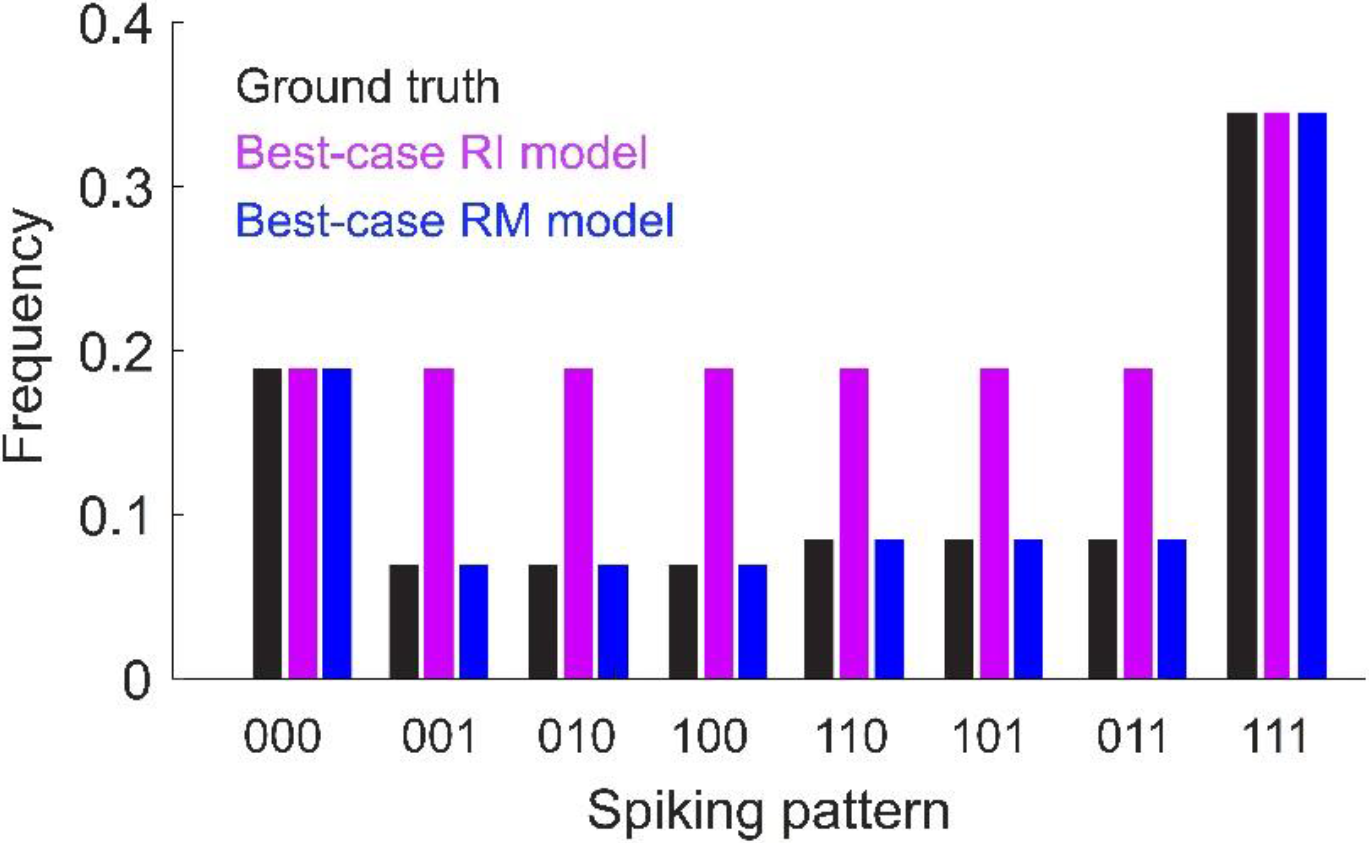
Toy model of *N* = 3 neurons with only first-and second-order interactions. Ground-truth probabilities are shown for each spiking pattern (black). Also shown are the frequencies predicted by the best-case (i.e., assuming infinite data and fitting time) Reliable Interaction (RI, magenta) and Reliable Moment (RM, blue) models (assuming a threshold of 0.1). Under these assumptions, the RM model would fit the ground-truth frequencies exactly. The RI model exactly fits the frequencies for spiking patterns above threshold, but is inaccurate for rare patterns. Note that the RI model cannot be normalized because the fitted partition function does not match fitted interaction terms (see main text and Appendix A for a detailed explanation). Model parameters: *α* = 1, *β* = 1.2 (see Equation (3)).

On the other hand, because it falls in the class of maximum entropy distributions, the RM model is guaranteed to converge to the ground-truth solution under the following assumptions: first, assuming that all interaction terms in the ground-truth model are incorporated into the RM model; second, assuming infinite data; and finally, assuming infinite time and a convex iterative fitting procedure such as Iterative Scaling [24]. For this toy example, this means that the “best case” RM model given by Equation (5) will converge to the ground-truth distribution (Equation (3)). However, note that this is not necessarily the case due to sampling noise, unidentified interaction terms, and the necessity for approximate methods due to time limitations. In the latter case, we advocate the use of the approximate MPF learning algorithm as a more practical option than Iterative Scaling, but this choice introduces some error into the fitted model. Approximate methods are also useful for calculating the partition function. While the partition function can be calculated exactly by brute-force summing all 2^N^ unnormalized probabilities, this can become prohibitively slow for large populations. We instead approximate the partition function; e.g., by the Good−Turing estimate [23]. Another alternative is to use Gibbs sampling [25] to generate spiking patterns from the inferred interaction parameters, then use the RI estimate of the partition function as the inverse probability of the non-spiking state in the Gibbs sampled “data”. Regardless of which of these methods is used, our toy example shows the fundamental differences between the RM and RI models, namely, that the RM model can in principle be normalized without disrupting its predictions of spike pattern probabilities.

### 2.3. The RM Model Infers Fewer Strong Spurious Higher-Order Interactions

Using this toy model, we have demonstrated that the RM model may be: (1) less likely to infer spurious higher-order interactions, and (2) better able to predict the frequencies of new spiking patterns. Do these improvements hold for more realistic population spiking statistics? To test this, we modeled populations of *N* = 20 neurons using pairwise maximum entropy models. Specifying the desired statistics of a maximum entropy model is a notoriously difficult inverse problem. We therefore tuned the ground-truth interaction parameters to generally give low firing rates (Figure 2a, mean ± std, 3.3 ± 1.9 Hz) and a broad distribution of correlations (Figure 2b, 0.01 ± 0.05; see Methods). However, we will subsequently test the ability of the RM model to fit a class of models for which we can directly prescribe cortical-like distributions of firing rates and spiking correlations (see Section 2.5).

**Figure 2.**
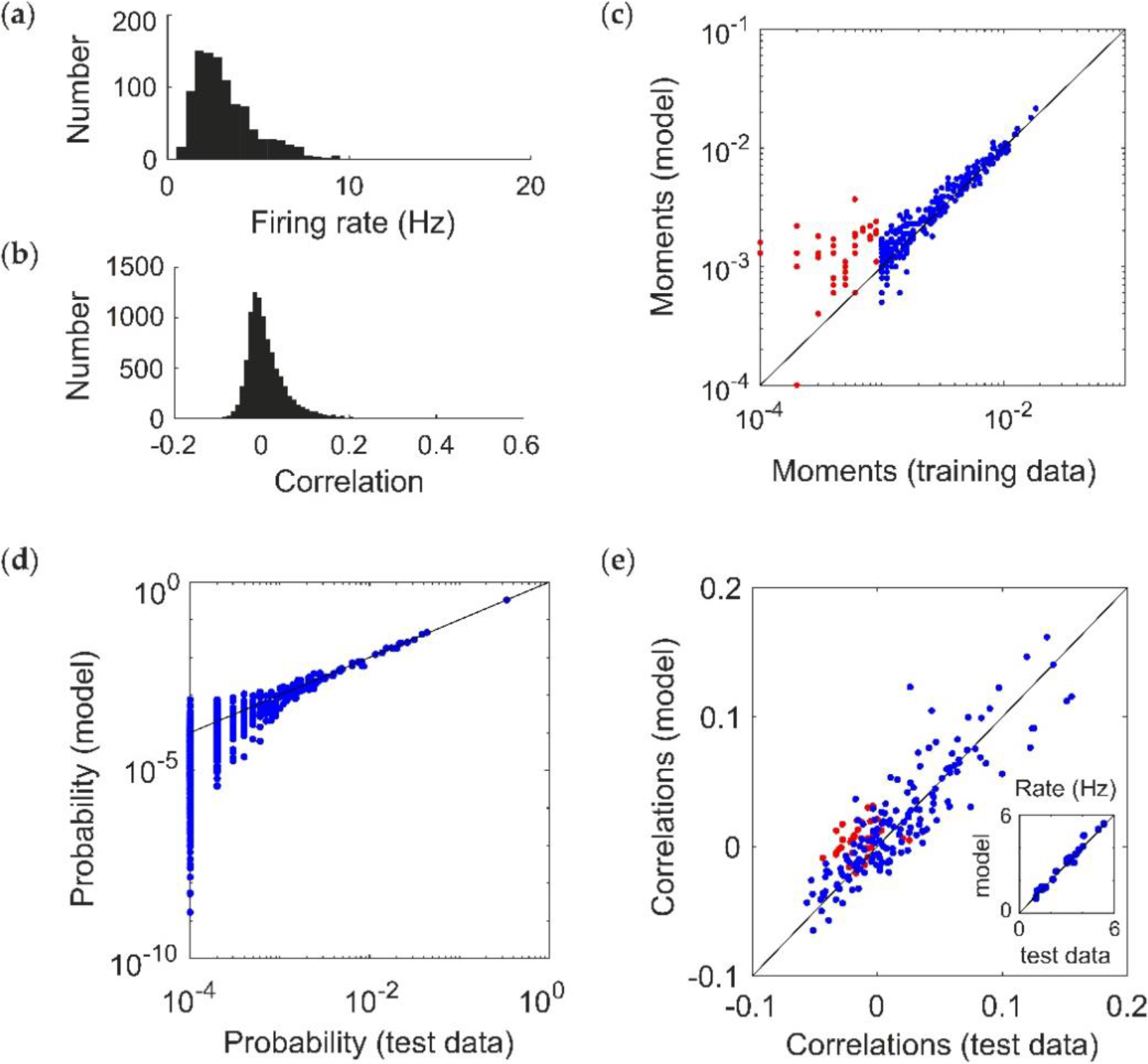
Fitting a ground-truth pairwise maximum entropy model (*N* = 20). (**a**,**b**) Distribution of (**a**) firing rates (assuming a time window of 20 ms) and (**b**) pairwise correlation coefficients generated by the ground truth models. (**c-e**) Example of Reliable Moment (RM) model fit to 200 s of a simulated pairwise ground truth model (*p*_*min*_ = 10^−3^). In this example, the RM model identified all 20 units, 154 pairs, 103 triplets, and 5 quadruplets. (**c**) Uncentered sample moments in the fitted RM model plotted against the empirical sample moments (estimated from training data) to show quality of model fit. Blue indicates all moments (single, pairwise, and higher-order) that were identified by the RM model. For comparison, red indicates the 36 pairs that were not identified by the RM model (and hence not fitted).(**d**) Cross-validated RM model probabilities versus ground-truth probability (i.e., estimated from held-out “test” data), for an example ground-truth model. Each point represents a different spiking pattern. (**e**) RM model correlations plotted against cross-validated empirical correlations (i.e., sample correlations plotted against empirical sample correlations from test data). Again, red points indicate pairs whose corresponding interaction terms were not identified. Inset shows the same for firing rates.

We generated population spiking patterns under the resulting distribution using Gibbs sampling (equivalent to 200 s worth of data) [22,25]. Figure 2c shows the fitted moments of a RM model for an example simulated population dataset (*p*_*min*_ = 10^−3^). This choice of threshold parameter identifies all 20 units, and 154 pairs (out of 190 possible) as having moments above threshold, which are fitted via MPF learning to reproduce the sample moments from the training data (blue; for comparison, the 36 pairs that were not included in the fitting are shown in red). The model is able to reproduce the probability distribution of spiking patterns in a “test” dataset that was not used to fit the model (Figure 1d), as well as the firing rates and correlations (Figure 1e; including the pairs that were not explicitly used for fitting, in red). This choice of model also identifies 108 higher-order moments (103 triplets and 5 quadruplets) as being above threshold. Since the ground truth model is pairwise, ideally their interaction parameters should be zero after fitting. Because of sampling noise in the data, as well as idiosyncrasies of MPF learning (see Discussion), they are nonzero but small on average (magnitude 0.235 ± 0.231).

How does this compare to the RI model? We next systematically tested whether the RM and RI models infer spurious higher-order interactions by simulating 50 random pairwise populations (using the same firing rates and correlations given by the distributions in Figure 2a,b). For each ground-truth model, we fit 20 RM and RI models with varying thresholds (see Methods), and compared the magnitudes of the higher-order interaction parameters. We found that the fitted higher-order interaction terms were smaller for the RM model than the RI model, regardless of the number of inferred parameters (Figure 3). This was true even when correcting for potential differences in the fitted lower-order interaction parameters (see Appendix B). Moreover, for the RM model, the average magnitude of the higher-order interaction terms was nonzero, but small and constant across different thresholds; whereas for the RI model, they increased in both magnitude and in variance. When a sparse subset of triplet interaction terms is added to the ground-truth model, the RM model is also better able to fit the corresponding interaction parameters (see Appendix C). These results reinforce the intuition we developed previously with the toy model (Figure 1) that the RM model finds fewer strong, spurious higher-order interactions, and is better able to fit existing higher-order interactions.

**Figure 3.**
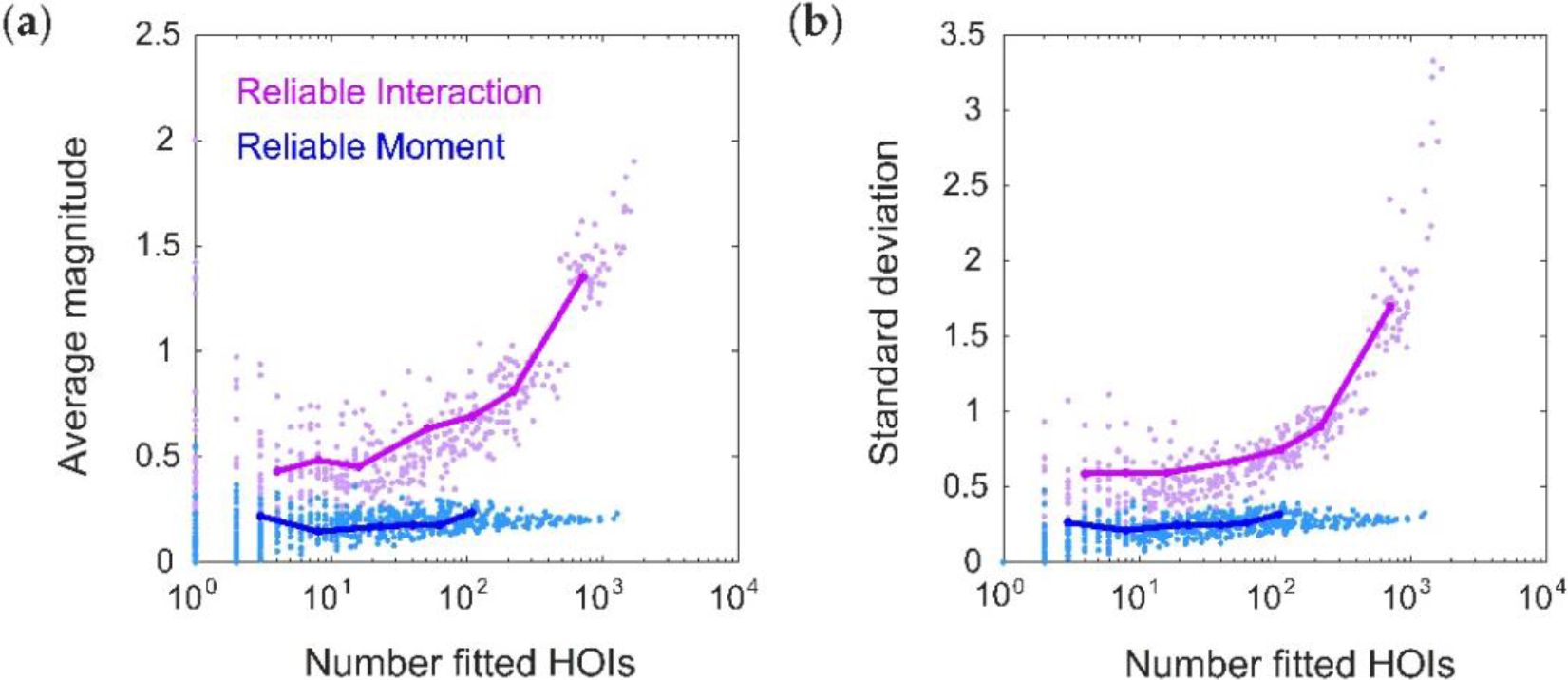
The Reliable Moment (RM) infers fewer strong, spurious higher-order interactions. (**a**) Average magnitude of all fitted higher-order interaction parameters as a function of the number of fitted higher-order interactions, shown for both the Reliable Interaction (RI; magenta) and RM (blue) models. Note that all higher-order interactions should have magnitude 0. Points represent 50 random ground-truth models (i.e., random interaction parameters), each of which is fitted 20 times with varying threshold parameters (see Methods). Solid lines indicate the RM and RI fits to a specific example ground-truth model. (**b**) Same as (a) but for standard deviation.

### 2.4. The RM Model Fits Rare Spiking Patterns

Our toy model also predicted that, while the RI model is very accurate at capturing the frequencies of commonly occurring spiking patterns, it is unable to predict the probabilities of rare patterns. This could be a strong limitation for large population recordings, as the number of previously-unseen spiking patterns grows as O(2^N^) assuming fixed recording lengths. We therefore tested this effect by generating a new testing dataset for each ground-truth model, and separating it into “old” spiking patterns (those that also occurred within the training dataset) and “new “ spiking patterns (those that only occurred within the test dataset). In order to compare the RM and RI models, we must specify which threshold values to use for each model. Since the RM and RI threshold use different “units” (i.e., the RI threshold is based on the frequencies population spiking patterns, and the RM threshold is based on marginal probabilities or moments), it is difficult to directly compare them. For a fair comparison of the model fits, it is therefore necessary to compare models that have the same number of fitted interaction parameters. Otherwise, any difference in model performance might be attributed to a model having more parameters to fit. We therefore first chose the threshold parameters in this example so that the RM and RI models have exactly the same number of fitted interaction parameters (in this case, 395). Figure 4a shows an example of model vs. empirical frequencies (calculated from held-out test data) for old spiking patterns.

**Figure 4.**
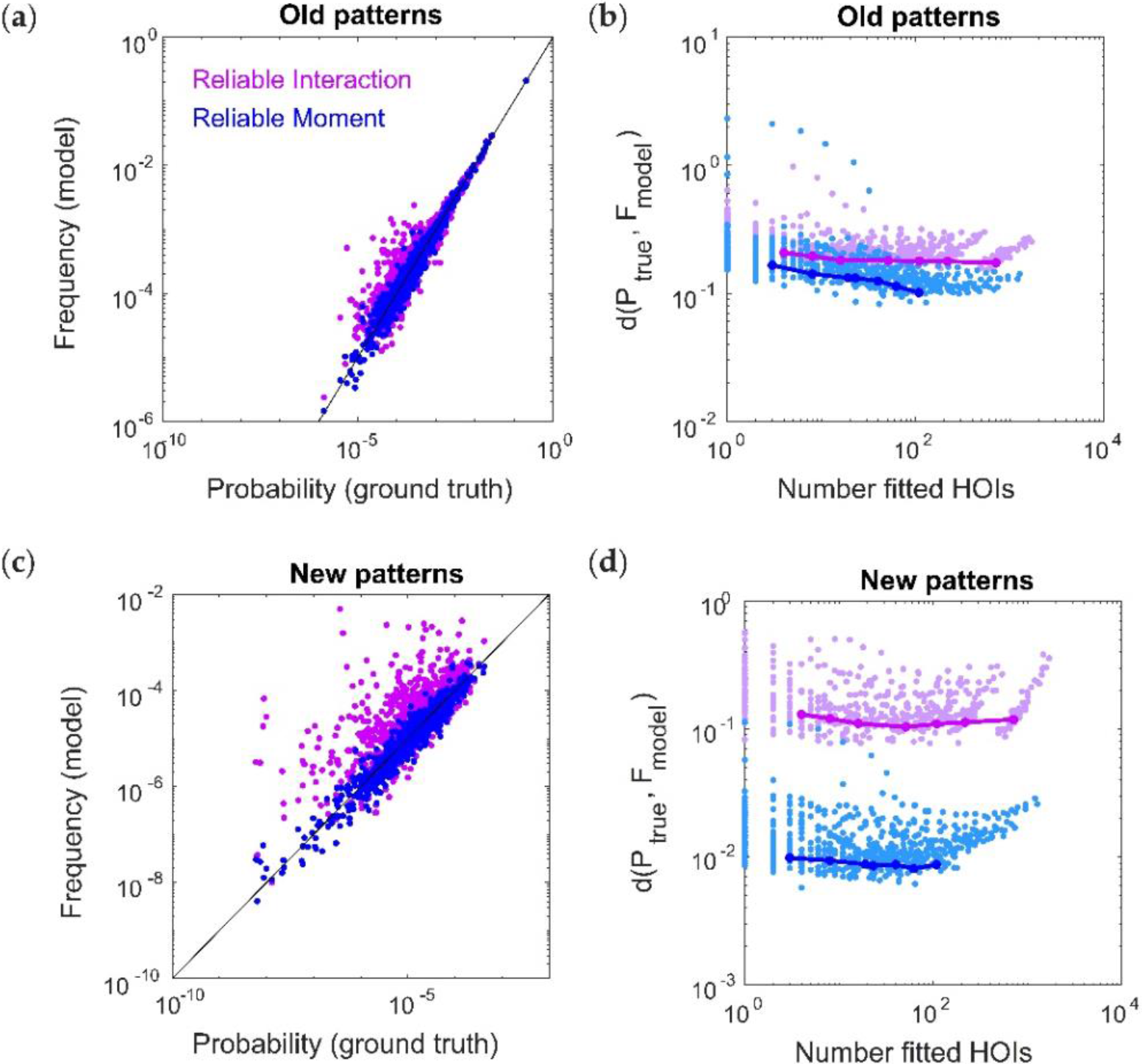
The Reliable Moment (RM) model is able to predict the probabilities of new spiking patterns. (**a**) Reliable interaction (RI; magenta) model frequencies and RM (blue) model probabilities of previously observed spiking patterns plotted against ground-truth probability, for an example ground-truth model. Each point represents a different “old” spiking pattern (i.e., occurring within both test and training datasets). For a fair comparison, we chose an example in which the RM and RI models had the same number of fitted interaction parameters (in this case, 395). (**b**) Dissimilarity (see Methods) between ground-truth distribution and model distribution of spiking patterns over different numbers of fitted higher-order interactions. Points represent 50 random ground-truth models (i.e., random interaction parameters), each of which is fitted 20 times with varying threshold parameters. Solid lines indicate the RM and RI fits to a specific example ground-truth model. (**c**,**d**) Same as (**a**,**b**) for new spiking patterns (i.e., those observed in the test data but not observed in the training data).

Because the RI model is unnormalized, we cannot use the Kullback−Leibler divergence. Instead, we calculated the dissimilarity between the distributions using the weighted average of the magnitude of the log-likelihood (see Methods, [1]). RM and RI model performances were comparable across different ground-truth populations and different threshold parameters (Figure 4b). However, the RI model was much less accurate for predicting the frequencies of new spiking patterns (Figure 4c,d). As discussed for the toy model, this is because the RI fitting procedure is only able to capture data that was used for fitting, which precludes new spiking patterns. Therefore, in both the toy model and the more realistic case here, the RM model is better able to predict the frequencies of the many unobserved spiking patterns that inevitably occur in large array recordings.

### 2.5. Fitting a Model with Cortical-Like Statistics and Dense Higher-Order Correlations

Thus far we have focused on fitting data generated by ground-truth pairwise maximum entropy models. Therefore, we now test the performance of the RM model on the Dichotomized Gaussian (DG) model, which simulates population spiking activity by generating multivariate Gaussian samples (representing correlated inputs to the population) and thresholding them [26]. The DG model generates dense higher-order statistics and can reproduce higher-order correlations observed in cortical data [9]. Unlike maximum entropy models, we can directly specify the firing rates for the DG model in order to generate cortical-like statistics. We chose log-normal, low-rate (mean 4 Hz) firing rate distributions [27,28] (Figure 5a), and normally distributed (mean 0.1) pairwise correlations [29] (Figure 5b; see Methods). We next compared the ability of the RM and RI models to fit the DG model spike patterns by comparing the dissimilarity between the model frequencies and the empirical probabilities from a held-out test dataset. The RM model was able to fit the DG patterns well, with the classic U-shaped curve with the number of parameters, whereas the RI model had an oscillatory shape (Figure 5c). The oscillations occur due to instabilities in the model’s ability to fit rare spiking patterns. To see this, Figure 5d shows an example of cross-validated model vs. empirical frequencies of spiking patterns that occur more than once in the test dataset. This is analogous to comparing the performance of the models for old spiking patterns (as in Figure 4a,b). For a fair comparison, we chose this example so that the RM and RI models had the same number of fitted interaction parameters (in this case, 239). Both models describe the data well, with the RI model performing slightly better because of its more accurate fit to the most common (quiescent) spiking patterns. However, when considering all the spiking patterns that occur in the test dataset, the RI model produces incoherent values for rare spiking patterns (i.e., those that only occur once, analogous to the “new” spiking patterns in Figure 4c,d), with frequencies often far surpassing 1 (Figure 5d, inset). Finally, note that an advantage of the RI model is that its fitting procedure is essentially instantaneous (Figure 5e). We therefore conclude that the RI model is a highly efficient method for capturing the frequencies of observed spiking patterns with relatively few parameters, but is unstable for predicting previously-unseen spiking patterns.

**Figure 5.**
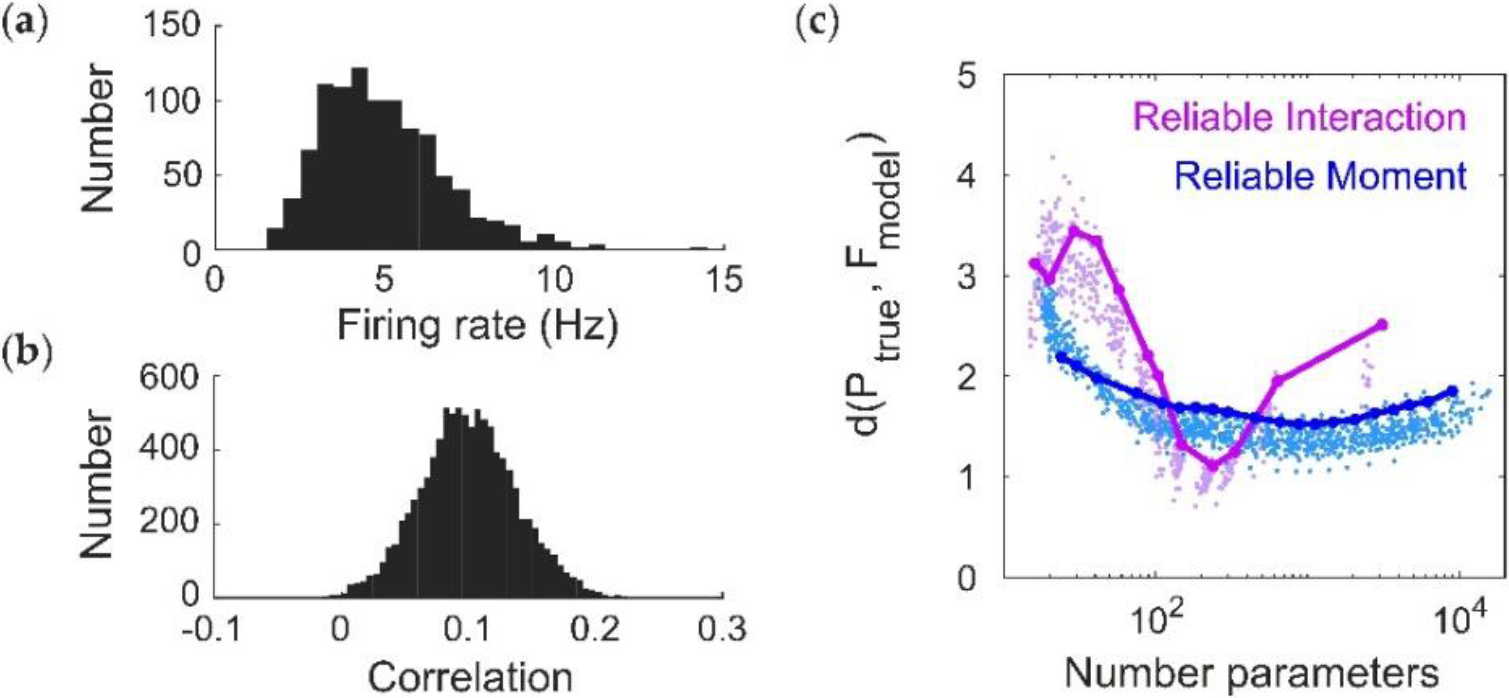
Fitting a Dichotomized Gaussian model with cortical-like statistics (*N* = 20). (**a**,**b**) Distribution of (**a**) firing rates (assuming a time window of 20 ms) and (**b**) pairwise correlation coefficients generated by the model. The Dichotomized Gaussian model is known to generate dense higher-order correlations [9,26]. (**c**) Cross-validated dissimilarity between the empirical and model distributions, for both Reliable Interaction (RI; magenta) and Reliable Moment (RM; blue) models. Points represent 50 random ground-truth models (i.e., random interaction parameters), each of which is fitted 20 times with varying threshold parameters. Solid lines indicate the RM and RI fits to a specific example ground-truth model. (**d**) Cross-validated model frequencies versus empirical probability, for an example ground-truth model. Each point represents a different spiking pattern. Only patterns occurring at least twice in the dataset are shown. Inset shows same plot, including spiking patterns that only occur once. For a fair comparison, we chose an example in which the RM and RI models had the same number of fitted interaction parameters (in this case, 239). (**e**) Time required for fitting RM and RI models.

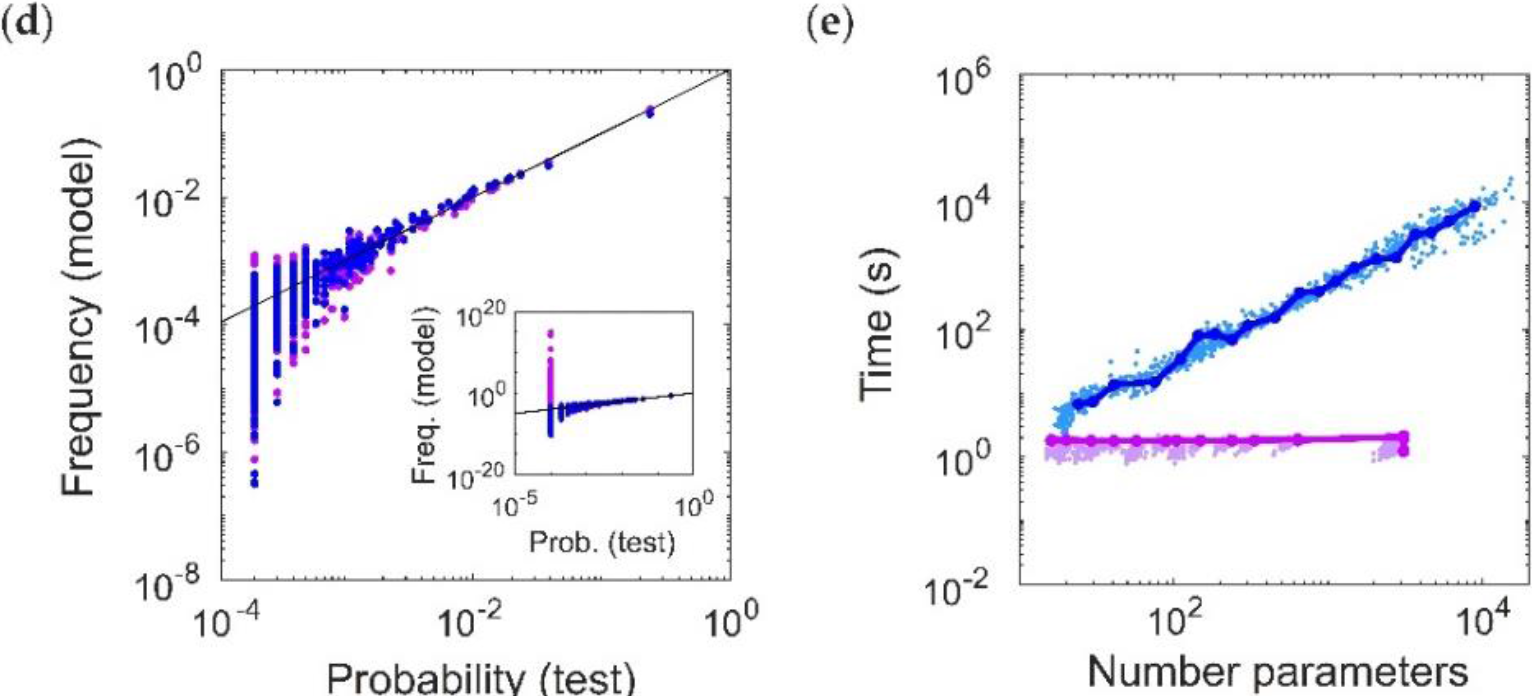

## 3. Discussion

We developed the Reliable Moment (RM) model, a novel class of maximum entropy model for adaptively identifying and fitting pairwise and higher-order interactions to neural data. To do this, we extended a previous model [1] by making two key modifications in the fitting criterion and the fitting procedure. First, we include spiking features whose corresponding uncentered moments are above a threshold value. This threshold need not be arbitrary, as it can be used to bound the confidence interval of the relative error (Equation (2)) [14]. Second, we take advantage of recent fast parameter fitting techniques [22], which results in a normalized probability distribution. We show that the RM model is able to fit population spike trains with cortical-like statistics, while inferring fewer strong, spurious higher-order correlations, and more accurately predicting the frequencies of rare spiking patterns.

We extended the intuition of the Reliable Interaction (RI) model [1] by determining which spiking features were most “reliable” as a criterion for inclusion in the model. However, our modifications confer several benefits. First, the RM model is normalized. While this does not necessarily affect the ability of a model to fit spiking pattern frequencies, it means that certain quantities that depend on the full distribution, such as mutual information or specific heat, can be applied to the RM model (although the RI model can be used to decode spiking patterns, as in [1]). This allows the RM model to be used for analyzing population coding or Bayesian inference, or for measuring signatures of criticality [17]. Second, the RM model is better able to predict the frequencies of previously-unseen spiking patterns. This is important for neural data, as the number of unseen spiking patterns increases significantly for large-scale population recordings. On the other hand, as a result of its fitting method, the RI model can be unstable for rare patterns (Figure 5d, inset; although it is able to predict the frequencies of common patterns well). Third, the RM model is less likely to find spurious higher-order interactions in a pairwise model, as compared to the RI model. This is because the hierarchical structure of the uncentered moments guarantees that no higher-order spiking feature can be fitted without also fitting all of its lower-order feature subsets. Finally, the main disadvantage of the RM model is that it is much slower to fit than the RI model, even using Minimum Probability Flow learning [22]. Therefore, the RM model performs better for determining the higher-order statistical structure of the data or predicting the frequencies of new patterns, while the RI model performs better as a fast method for fitting commonly occurring spiking patterns.

Several variants on the RM model are possible. While we chose to use MPF learning due to its speed, there are many alternative methods that are available [24,30–32]. In particular, classic Iterative Scaling [24] finds the interaction parameters that maximize the log-likelihood of the data. This is equivalent to minimizing the Kullback−Leibler divergence between the data and the model, which can be shown to be a convex problem. However, it can be prohibitively slow even for reasonably sized populations. On the other hand, MPF defines a dynamical system that would transform the empirical distribution into the model distribution, then minimizes the Kullback−Leibler divergence between the empirical distribution and the distribution established by this flow (after an infinitesimal time) [22]. While there is no guarantee on convexity, MPF in general works very well in practice (see Figure 2) and is much faster. Another possibility is to add a regularization term to the cost function during fitting to ensure sparse interaction parameters. Moreover, there is some flexibility in choosing the threshold parameter. Here, we advocated determining the threshold parameter to bound the error of the moments (Equation (2)). An alternative option would be to use the Akaike Information Criterion to determine the threshold that results in the optimal number of interaction parameters [33]; however, this would require multiple model fittings for validation. The criterion for inclusion of specific interactions may also be modified, for instance, by requiring that fitted interaction parameters have moments exceeding a threshold based on the empirical values. For each of these variants, the RM model extends the ideas behind the RI model by fitting a sparse subset of the most “relevant” higher-order interactions, while ensuring that the corresponding lower-order interactions are also fit.

We focused on capturing stationary correlations in neural data, while neglecting temporal dependencies between neurons. In principle, temporal correlations could be incorporated into the RM model, and into the maximum entropy models more generally, by fitting concatenated spatiotemporal spiking patterns [34–36]. This dramatically increases the cost of fitting, although emerging techniques are making this problem more tractable [37,38]. Another, more widely-used approach to fitting spatiotemporal models of neural spiking is the generalized linear model (GLM) [39,40]. GLMs and Maximum entropy models are not mutually exclusive; indeed, hybrid approaches are possible [41], and maximum entropy models can be interpreted in a linear-nonlinear framework [15,19]. Future work could incorporate higher-order moments into the GLM framework, as has been done for common inputs [42]; indeed, there is a long history of moment-based methods for point process models that could be drawn upon [43–46]. Such an advance could provide a powerful tool for fitting higher-order spatiotemporal statistics to neural circuits, and help to illuminate the structure of collective neural activity.

## 4. Materials and Methods

### 4.1. Ground Truth Models

We simulated ground-truth pairwise maximum entropy models for *N* = 20 neurons. Throughout, we assumed time bins of 20 ms for the spiking patterns. To test the performance of the Reliable Moment (RM) model on data without any higher-order interactions, we first assumed a pairwise maximum entropy distribution of spiking patterns with random, normally distributed first and second-order interaction parameters: *h*_*i*_~𝒩(3,0.25), 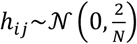. The metaparameters for the distributions were tuned to give low average firing rates and a broad distribution of correlations (Figure 2a). To calculate probabilities from the ground-truth model, we either calculate the empirical frequency of spiking patterns (‘empirical probability’, Figure 2d) or else we calculate the exact probability from model parameters (“ground-truth probability”, Figure 4). The latter requires an expensive calculation of the partition function.

For cortical-like models with higher-order statistics, we used a technique based on the Dichotomized Gaussian (DG) model [47] to generate spike trains with specified firing rates and spiking correlations. In this case, we drew firing rates from a lognormal distribution with a mean of 4 Hz and standard deviation of 2 Hz, and correlations were normally distributed ρ_*ij*_~𝒩(0.1,0.05). In this case, all probabilities are calculated based on empirical frequency (Figure 5).

### 4.2. Identification of Reliable Moments

To fit the RM model, we must first identify which moments are greater than *p*_*min*_. This process can be made efficient by taking advantage of the hierarchical arrangement of moments. We first find the set of neurons whose mean firing rates in the training data are greater than threshold:

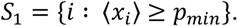

*S*_1_ is the set of first-order interaction parameters. Similarly, the set of *k*th-order interaction parameters is given by:

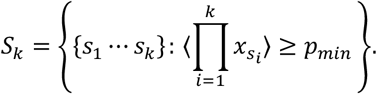

The RM model fits the interactions corresponding to all elements in *S*_*k*_, *k* = l,…,*N*. Enumerating all *S*_*k*_ can be computationally expensive as the number of possible interactions increases as O(N^k^). Because of the hierarchy of moments, this search can be expedited by only considering the *k*th-order subsets {*s*_1_ …*s*_*k*_} for which all of their (*k* − 1)th-order subsets are elements of *S*_*k*−1_. This determines whether the corresponding moment is above threshold. This step is performed iteratively until *S*_*k*_ = ø.

### 4.3. Model Fitting and Sampling

We fit the interaction parameters for the RM model using Minimum Probability Flow learning [22], which we adapted to accommodate arbitrary spiking interactions. After fitting the model, we used the Good-Turing estimate [23] to estimate the partition function empirically. For each ground-truth model, we fit 20 RM models with threshold parameters varying from *p*_*min*_ = 0.05 to *p*_*min*_ = 0.001. Because MPF is not convex (and therefore not guaranteed to converge), it is important to check that the model correlations reproduce the data correlations. To do this, we calculate sample correlations via Gibbs sampling.

The Reliable Interaction (RI) models were fit using the procedure described in [1]. Because spiking pattern frequencies are smaller than the marginal frequencies, we used smaller thresholds for the RI model, ranging from *p*_*min*_ = 5 * 10^−3^ to 10^−5^, as these resulted in similar numbers of fitted parameters in the RM and RI models.

### 4.4. Dissimilarity Between Empirical Data and Models

Since the RI model is not normalized, the Kullback−Leibler divergence returns incongruent (negative) values. We therefore follow [1] in measuring the dissimilarity between the ground-truth and the model frequencies as:

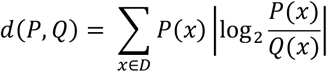

where 𝒟 is the set of all observed spiking patterns in the test data (however, in contrast to [1], we do not exclude spiking patterns that only occurred once).

### 4.5. Code Availability

All relevant code will be made available on GitHub.

## Author Contributions

N.A.C.G., J.Z., and E.S.B. conceived and designed the experiments; N.A.C.G. performed the computational experiments and analysis; N.A.C.G., J.Z., and E.S.B. wrote the paper.

## Acknowledgments

J.Z. gratefully acknowledges the following funding: Google Faculty Research Award, Sloan Research Fellowship, and Canadian Institute for Advanced Research (CIFAR) Azrieli Global Scholar Award. This work was also supported by an NSF Grants 1514743 and 1056125 to E.S.B.

## Appendix A

Here we demonstrate why the RI model cannot be normalized, using an intuitive example that can only be described in the maximum entropy formulation in the limit that the interaction parameters → −∞. However, note that the RI model for the toy example discussed in the main text (described by Equations (3) and (4)) is also unnormalizable in its exact form.

Consider *N* neurons that never spike: then, *P*(00…0) = 1, and zero for all other spiking patterns. Informally, this distribution can be described by the limit of the following first-order maximum entropy model:

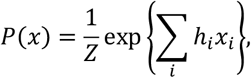

as *h*_*i*_ → − ∞. Under the RI model, the partition function is estimated as 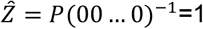. Since this is the only occurring pattern, all interactions are set to 0. As a result, the frequency of any spiking pattern is:

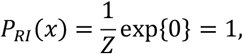

so that the frequencies sum to 2^N^. Although the RI model accurately (in this example, perfectly) fits the most common spiking pattern (silence), it is unnormalized. Furthermore, naive renormalization would result in a probability distribution that is inaccurate for every spiking pattern. This dilemma occurs because 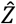 is an accurate (in this example, perfect) estimate of the partition function of the underlying distribution, but not for the model defined by the interaction parameters identified by the RI model.

## Appendix B

We have shown that the RM model predicts infers smaller higher-order interaction parameters in a ground-truth pairwise model than the RI model. In principle, this fact could be due to changes in the lower-order terms. In other words, it is possible that the RI higher-order interaction terms are larger in absolute magnitude, but not in relative magnitude as compared to the e.g., pairwise terms. We therefore quantified the average magnitude of fitted higher-order interaction parameters, normalized by the average magnitude of all fitted pairwise terms. However, we still found that the normalized higher-order interaction terms were larger in the RI model than in the RM model (Figure A1a). This is due to the fact that the pairwise interaction terms were similar between the RM and RI models (Figure A1b). Therefore, we conclude that the RM model infers fewer strong, spurious higher-order interactions, even when controlling for differences in the lower-order terms.

**Figure A1.**
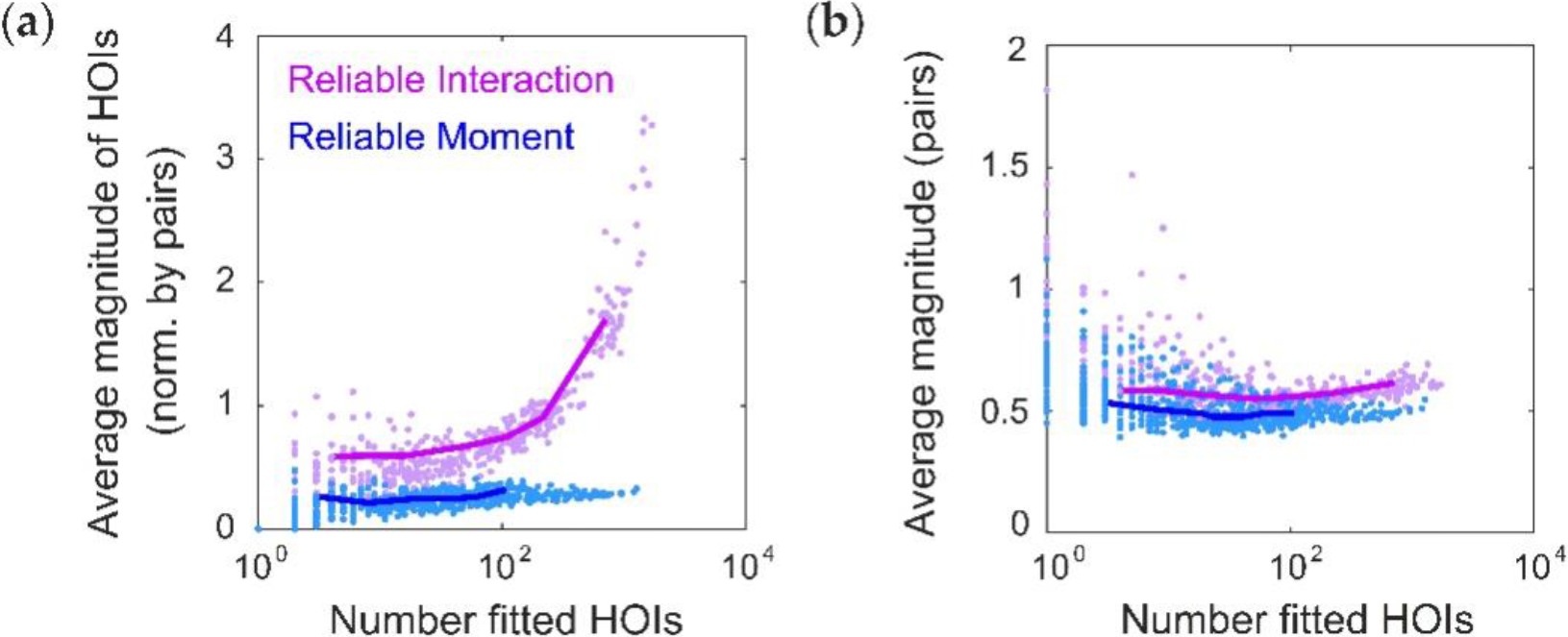
The Reliable Moment (RM) model infers smaller parameters for spurious higher-order interaction terms, relative to lower-order terms. (**a**) Average magnitude of all fitted higher-order interaction parameters normalized by the average magnitude of all pairwise interaction parameters, shown for both the Reliable Interaction (RI; magenta) and RM (blue) models (cf. Figure 3a). (**b**) Average magnitude of all pairwise interaction parameters (RI, magenta; RM, blue).

## Appendix C

To test whether the RM model is better able to fit higher-order interactions than the RI model, we augmented the pairwise maximum entropy model with a sparse set of nonzero triplet interaction terms. Specifically, the lower-order terms were generated in the same manner as for the pairwise model (see Methods). Then, each of the 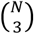 possible triplet terms was chosen with probability *p* = 0.05, and the corresponding interaction parameters for these triplets (referred to as the “ground-truth triplets”) were drawn from a standard normal distribution. We repeated the same fitting protocol as previously described for the pairwise maximum entropy model. The number of ground-truth triplets that were not identified by the RM model was slightly higher than the RI model (mean ± std, 53.00 ± 0.29, RM model; 46.00 ± 0.37, RI model). However, we found that the inferred interaction parameters for these ground-truth triplets are more accurate for the RM model than the RI model (Figure A2). Therefore, while the RM model misses slightly more ground-truth triplets than the RI model, it more accurately fits their interaction parameters.

**Figure A2.**
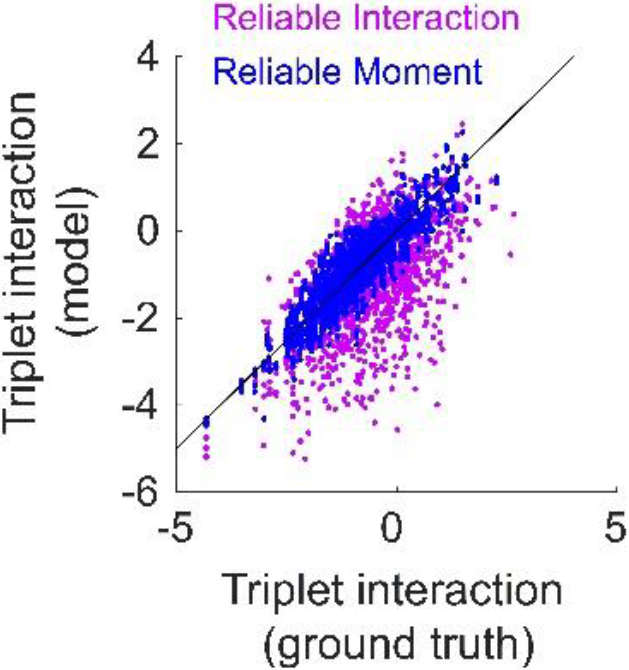
The Reliable Moment (RM) model more accurately fits higher-order interaction parameters of a maximum entropy ground-truth model incorporating a sparse subset of triplet terms. Fitted interaction parameters for ground-truth triplets inferred by the RM model (blue) and the Reliable Interaction model (RI, magenta), plotted against the ground-truth values of the interaction parameters.

